# Serinc5 restricts HIV membrane fusion by altering lipid order and heterogeneity in the viral membrane

**DOI:** 10.1101/2022.08.25.505328

**Authors:** Amanda E. Ward, Daria Sokovikova, M. Neal Waxham, Frederick A. Heberle, Ilya Levental, Kandice R. Levental, Volker Kiessling, Judith M. White, Lukas K. Tamm

## Abstract

The host restriction factor, Serinc5, incorporates into budding HIV particles and inhibits their infection by an incompletely understood mechanism. We have previously reported that Serinc5 but not its paralogue, Serinc2, blocks HIV cell entry by membrane fusion, specifically by inhibiting fusion pore formation and dilation. A compelling body of work also suggests Serinc5 may alter the conformation and clustering of the HIV fusion protein, Env. To contribute an additional perspective to the developing model of Serinc5 restriction, we assessed Serinc2 and Serinc5’s effects on HIV pseudoviral membranes. Using fluorescence lifetime imaging with an order sensitive dye, FLIPPER-TR, and by measuring pseudoviral membrane thickness via cryo electron microscopy (cryoEM), Serinc5 was found to increase membrane heterogeneity, skewing the distribution towards a larger fraction of the viral membrane in an ordered phase. We also directly observed for the first time the coexistence of membrane domains within individual viral membrane envelopes. Using a TIRF-based single particle fusion assay, we found that incorporation of exogenous phosphatidylethanolamine (PE) into the viral membrane rescued HIV pseudovirus fusion from restriction by Serinc5, which was accompanied by decreased membrane heterogeneity and order. This effect was specific for PE and did not depend on acyl chain length or saturation. Together, these data suggest that Serinc5 alters multiple interrelated properties of the viral membrane—lipid chain order, rigidity, line tension, and lateral pressure—which decrease accessibility of fusion intermediates and disfavor completion of fusion. These biophysical insights into Serinc5 restriction of HIV infectivity could contribute to the development of novel antivirals that exploit the same weaknesses of HIV and potentially other enveloped viruses.

## Introduction

Serinc5 is a recently described host restriction factor that incorporates into budding HIV particles and inhibits infection at the cell entry step (Rosa et al., 2015; Usami et al., 2015). While it is known to inhibit multiple steps of membrane fusion—hemifusion, fusion pore opening, and fusion pore dilation (Sood et al., 2017; Ward et al., 2020)—the exact mechanism by which it does so remains uncertain. Many studies have focused on the interaction between Serinc5 and the HIV fusion protein, Env, demonstrating that Serinc5 alters Env conformation (Schulte et al., 2018; Staropoli et al., 2019), antibody binding (Beitari et al., 2017; Sood et al., 2017) and clustering (Chen et al., 2020). However, alterations in Env cannot fully explain the observed fusion defects induced by Serinc5 incorporation. In current models of viral membrane fusion, progression through receptor binding, hemifusion, and early fusion pore formation is driven by rearrangements of the viral fusion protein and membrane deformation (Harrison, 2015; White and Whittaker, 2016). However, the penultimate step, fusion pore dilation, is driven by membrane tension and curvature (Kozlov and Chernomordik, 2015). Disruption of fusion pore dilation, as we previously observed with Serinc5-containing HIV pseudoviruses (Ward et al., 2020), suggests that Serinc5 may alter fluid mechanical properties of the viral membrane. Increased viral membrane stiffness would require a larger input of energy to deform the membrane into highly curved intermediates like hemifusion stalks (Gracià et al., 2010) and fusion pores (Chernomordik and Kozlov, 2008). Changing lipid biophysical properties of the viral membrane (often via cholesterol depletion) has been shown to reduce fusion of HIV and other enveloped viruses (Biswas et al., 2008; Chlanda et al., 2016; Domanska et al., 2013; Haldar et al., 2019; Kasson and Pande, 2007; Lee et al., 2021; Sun and Whittaker, 2003; Wudiri et al., 2017; Yang et al., 2017, 2016b; Zawada et al., 2016).

While it has already been shown that Serinc5 does not change the overall lipid composition of the viral membrane (Trautz et al., 2017), membrane organization could be altered by other means, thereby affecting local concentrations of lipids. Model membranes of similar composition to the HIV membrane were shown to support liquid-liquid phase separation (Brügger et al., 2006; Huarte et al., 2016; Yang et al., 2015). Specifically, areas enriched in saturated phospholipids, cholesterol and sphingolipids form domains with a more ordered packing of lipid acyl chains that coexist with regions of unsaturated phospholipids, less cholesterol and fewer sphingolipids packed in a more disordered manner; these two phases are termed liquid-ordered (Lo) and liquid-disordered (Ld), respectively (Feigenson, 2006). As a consequence of ordered packing, membranes in an Lo phase have different lateral pressure profiles and are several angstroms thicker than membranes in an Ld phase (Levental et al., 2020). Lipid order and packing parameters can be reported by specific fluorescent membrane probes (Ashdown and Owen, 2015; Colom et al., 2018; Niko et al., 2016), some of which are also suitable for imaging in cellular and viral membranes. More recently, heterogeneity of membrane thickness was directly observed by cryogenic electron microscopy (cryoEM) in model membranes composed of pure lipids as well as isolated plasma membrane vesicles (Cornell et al., 2020; Heberle et al., 2020). Additionally, the host plasma membrane, from which the HIV viral membrane is derived, has an asymmetric lipid distribution, in which the outer leaflet is enriched in sphingomyelin (SM) and phosphatidylcholine (PC), while phosphatidylethanolamine (PE) and phosphatidylserine (PS) are actively sequestered on the inner leaflet. PE and PS are only present in the outer leaflet of the plasma membrane in times of cellular stress (Lorent et al., 2020), but are detectable in the outer leaflet of the HIV envelope (Amara and Mercer, 2015; Callahan et al., 2003; Chua et al., 2019; Huarte et al., 2016) and thus steady-state asymmetry of plasma membrane lipids is assumed to be lost in viral particles.

Previously, we showed that the impediment to membrane fusion of Serinc5-containing HIV pseudoviruses is overcome by incorporation of the exogenous lipid Atto488-dimyristoyl PE (Atto488-DMPE) and the lipophilic antifungal drug amphotericin B, while fusion of HIV particles containing the non-restricting paralogue, Serinc2, is unaffected by these additions (Ward et al., 2020). These data suggest that Serinc5 restriction may be dependent on the lipid environment of the viral membrane. To address the hypothesis that Serinc5 alters lipid bilayer properties of the HIV envelope, we systematically investigated the effects of Serinc5 incorporation on membrane order of pseudoviral particles and the physical and chemical properties of lipids required to overcome restriction by Serinc5.

## Results

### Serinc5 increases lipid acyl chain order in the viral membrane

To examine the effects of Serinc incorporation on the order and tension of the HIV lipid bilayer, we labeled HIV pseudovirus particles with the fluorescent membrane dye FLIPPER-TR. While FLIPPER-TR has been described as a reporter of membrane tension, it also reports on acyl chain packing in model or biological membranes. Viral membranes lack the typical sources of tension in cellular membranes (hydrostatic, cytoskeleton interactions, and adhesion) although it is possible that the HIV matrix protein (MA) introduces tension in the viral membrane (Kozlov and Chernomordik, 2015). Thus, in the context of viral membranes, FLIPPER-TR may primarily function as a reporter of membrane order and elastic modulus, i.e., membrane biophysical properties that are all inter-related. This push-pull fluorescent probe changes wavelength (red-shifted excitation) and has a longer lifetime in membranes with increased lipid order (Colom et al., 2018; Dal Molin et al., 2015; Licari et al., 2020). FLIPPER-TR has been used as a reporter of membrane order not only in lipid model membranes but also in a variety of biomembranes (Colom et al., 2018; Dal Molin et al., 2015; Goujon et al., 2019). To calibrate the FLIPPER-TR method for application to pseudoviruses, which appear as diffraction limited spots by fluorescence microscopy, we first measured the fluorescence lifetimes of FLIPPER-TR-stained large unilamellar vesicles (LUVs) of comparable size to pseudoviruses (Fig. 1, left and Fig. S1) by fluorescence lifetime imaging microscopy (FLIM). LUVs were made of ternary lipid compositions previously reported to exist as all L_o_, all L_d_, or co-existing L_o_/L_d_ phases (De Almeida et al., 2003; Ionova et al., 2012). As expected, we observed longer, shorter, and intermediate average lifetimes for L_o_, L_d_, and L_o_/L_d_ LUVs, respectively (Fig. 1, left). The measured lifetimes of FLIPPER-TR in all LUVs studied here are shorter than previously reported lifetimes in membrane vesicles with a larger diameter, which may be due to altered pressure profiles and elastic moduli in vesicles with decreasing diameters (Lin et al., 2012; Lipowsky, 2022). The lifetimes of FLIPPER-TR in these LUVs composed of varying ratios of sphingomyelin (SM), cholesterol, and palmitoyl-oleoylphosphatidylcholine (POPC) were also comparable to LUVs with simpler compositions known to exist as all L_o_, all L_d_, or co-existing L_o_/L_d_ phases (Fig. S2).

**Fig. 1.**
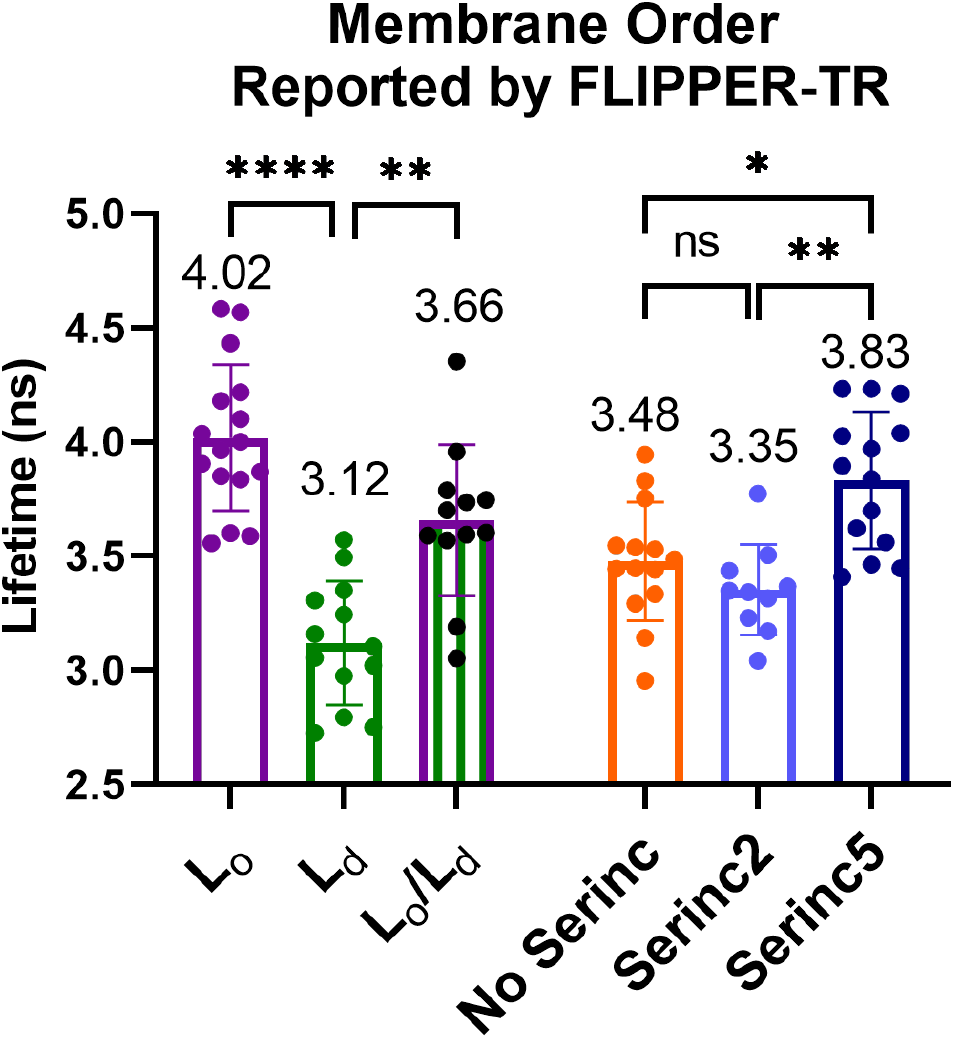
Fluorescence lifetimes of the lipid order sensitive membrane dye, FLIPPER-TR, in liposomes of defined lipid composition and in HIV pseudovirus particles. Large unilamellar liposomes (LUVs) composed of ternary mixtures of the major lipid components of the HIV envelope in ratios known to exist solely in L_o_ (35/40/25, SM/chol/POPC), solely L_d_ (25/5/70, SM/chol/POPC), or as a mixture of L_o_ and L_d_ (40/20/40, SM/chol/POPC) phases were used as standards for comparison. LUVs and HIV pseudoviruses were stained with FLIPPER-TR, adhered to a coverslip, and imaged by fluorescence microscopy. The fluorescent puncta were segmented from background by intensity thresholding and the average lifetime of all particles within a field of view are plotted as a point with mean and SD of all data points of identically prepared samples. The mean lifetime in nanoseconds of all data points for a condition is written above each bar. Data are from at least three independent preparations of LUVs and pseudoviruses. Multiple comparisons by Dunnett’s T3 multiple comparisons test: **** p<0.0001, ** p<0.01, * p<0.05, ns not significant.

HIV pseudoviruses were stained with FLIPPER-TR and imaged in the same manner by fluorescence lifetime microscopy. The lifetimes of the dye in pseudovirus membranes without Serincs or with Serinc2 (Fig. 1, right, orange and light blue circles, respectively), were similar to those in LUVs with coexisting L_o_ and L_d_ phases whereas the lifetimes in pseudovirus membranes that incorporate Serinc5 were longer, indicating more ordered membranes (Fig. 1, right, dark blue circles). These data show that unrestricted HIV pseudoviral membranes exhibit lipid chain order intermediate between the L_o_ and L_d_ extremes of model membranes and further support the hypothesis that Serinc5, but not Serinc2, alters the organization of lipids within the viral membrane to become more ordered.

### Viral membrane thickness is heterogeneous and Serinc5 broadens the viral membrane thickness distribution

To examine the effects of Serinc5 incorporation on viral membranes at higher resolution, we turned to recently developed techniques for visualizing membrane heterogeneity by cryoEM (Heberle et al., 2020). Increased membrane order is usually accompanied by an increase in membrane thickness. However, it is not trivial to measure thickness in biological membrane samples, especially if the membrane itself contains coexisting areas of variable thickness. This problem can be overcome by applying cryoEM to LUVs and biologically derived membranes (Cornell et al., 2020; Heberle et al., 2020). To detect membrane thickness variation in the envelopes of HIV pseudovirus particles, we flash-froze particles that incorporated Serinc2, Serinc5, or no Serincs and imaged them by cryoEM (Fig. 2). The membrane envelopes of the particles are characterized by two lines of high density, or “troughs” (Fig. S4C). The trough-to-trough distance (DTT) was previously found to match most closely with the hydrophobic thickness of the bilayer when compared with small-angle X-ray scattering (Heberle et al., 2020), suggesting that the trough position corresponds roughly to the glycerol backbone region of the bilayer. DTT of the viral membrane in these projection images was measured in 5 nm segments as described previously (Heberle et al., 2020) and example micrographs with DTT heat maps are shown in Fig. 2A and Fig. S3. We also frequently observe a line of density under the thinner portions of the membrane at a distance that is consistent with the matrix (MA) protein of HIV (Fig. S4).

**Fig. 2.**
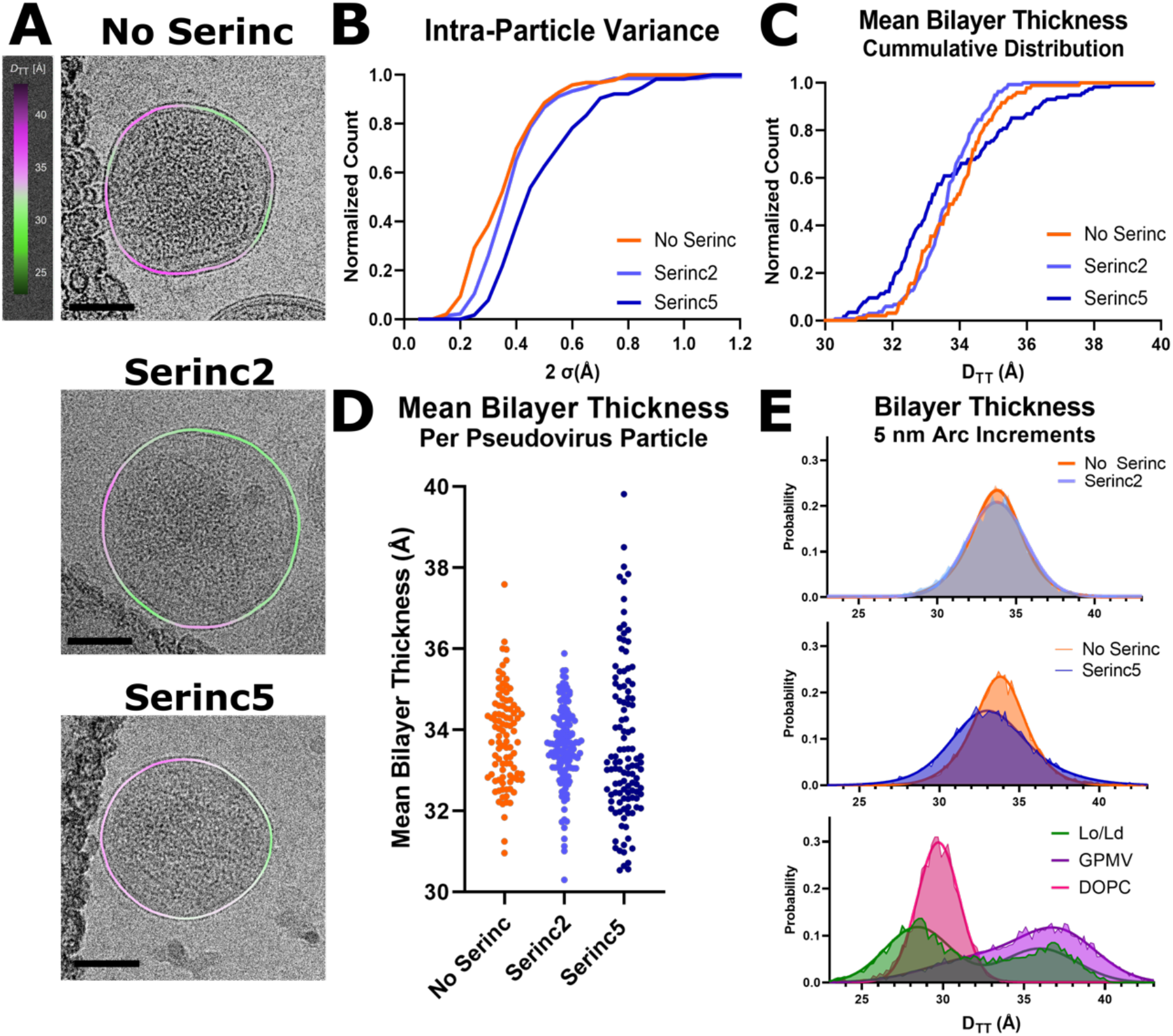
Serinc5 incorporation increases membrane heterogeneity and widens the thickness distribution of HIV pseudovirus particles as measured by cryoEM. (**A**) Example micrographs with membrane thickness (D_TT_) overlay of HIV pseudoviruses prepared without Serincs (No Serinc), with Serinc2, or with Serinc5. The measured distances between intensity troughs (D_TT_) for 5 nm segments of the membranes were plotted as a smoothed, colored overlay according to the scale on the left. Scale bars are 50 nm. (**B**) 95% confidence intervals of the mean bilayer thickness (D_TT_) of individual HIV pseudovirus particles plotted as a cumulative distribution function. Mean membrane thickness (D_TT_) of individual HIV pseudoviruses plotted as a cumulative distribution function (**C**) and as a bee swarm plot (**D**). Each point represents one pseudovirus particle. Data are combined from at least two independent preparations of pseudoviruses with over 100 viral particles analyzed for each condition. (**E**) Normalized distributions of membrane thickness (D_TT_) measurements of 5 nm segments of HIV pseudovirus, LUV, or GPMV membranes with double Gaussian fits. 1-phase LUVs were composed of DOPC/POPG (95/5 molar ratio) and 2-phase LUVs were composed of a 40/20/35/5 ratio of DPPC/chol/DOPC/POPG. LUV and GPMV distributions reproduced with permission from (Heberle et al., 2020).

The majority of particles from all three preparations show intra-particle variations in membrane thickness with contiguous areas of thicker or thinner membrane within any particle cross-section. Cumulative distribution plots of the standard deviation of membrane thickness within individual particles reveals substantially greater intra-particle thickness variance for Serinc5 particles compared to particles with Serinc2 or without Serincs (Fig. 2B). Averaging D_TT_ per particle, the mean bilayer thicknesses were similar for HIV particles with Serinc2 and without Serincs, but the distribution of thicknesses for Serinc5-containing pseudoviruses is much broader, with about 35% of all particles exhibiting thicker and about 65% thinner membranes than the average No Serinc or Serinc2 particle (Fig. 2C, D), reinforcing and expanding on the fluorescence lifetime data shown in Fig. 1. The broader distribution of membrane thicknesses in Serinc5 particles may be attributable to the known variability in Serinc5 incorporation into individual particles (Sood et al., 2017). The distribution of particle sizes is comparable for HIV pseudoviruses with and without Serincs and we detected no relationship between particle size and mean membrane thickness (Fig. S5). When the thicknesses of 5 nm segments of all analyzed pseudoviral membranes are plotted as probability histograms, the distributions of viral particles without Serincs and with Serinc2 are practically identical while the distribution from Serinc5-containing particles is broader, i.e. containing more regions of both thinner and thicker membrane areas (Fig. 2E). Comparing these pseudoviral distributions to previously published distributions of LUV and giant plasma membrane vesicle (GPMV) membranes (Heberle et al., 2020), all pseudoviral distributions show a single peak that is more broadly distributed than single phase LUVs, but still not as broadly distributed as GPMVs (Fig. 2E). While the pseudoviral distributions are centered between the L_d_ and L_o_ peaks of the two-phase LUVs, the GPMV distribution is skewed towards L_o_-like thicknesses. While membrane thicknesses in neither pseudoviral particles nor GPMVs were clearly bimodally distributed as they are in the two-phase LUVs, the broad distributions suggest a higher complexity in the lipid phase behavior of biological compared to model membranes. Broader distributions would also result if domain sizes were smaller and more numerous because out of plane domains of different thicknesses may appear as intermediate thickness in projection images. As domain size approaches the segment length of the analysis (5 nm), the probability increases that a segment contains densities of both phases, resulting in more segments with intermediate thickness and thus a broader unimodal distribution rather than a clearly bimodal distribution.

### Enrichment of the HIV pseudoviral membrane with PE reverses Serinc5’s fusion inhibition by decreasing membrane heterogeneity and order

We used a previously described total internal reflection fluorescence (TIRF) microscopy-based, single-particle fusion assay to examine the effects of enrichment of HIV pseudovirus particles with exogenous lipids (Fig. 3A). Briefly, binding of pseudovirus particles engineered to incorporate an mCherry content marker to a supported planar plasma membrane containing CD4 receptor and CCR5 co-receptor produces a sudden appearance of punctate fluorescence in the observed evanescent field. After remaining stably bound for several imaging frames, the particle either fuses, as reported by a decay of fluorescence as the mCherry content diffuses away over multiple frames of recorded video, or becomes unbound from the membrane, reported by a sudden drop in fluorescence intensity back to baseline over a single frame. We have previously established strong concordance between fusion of HIV pseudoviruses in this assay with *in vitro* infection efficiency, including particles with or without Serincs (Ward et al., 2020).

**Fig. 3.**
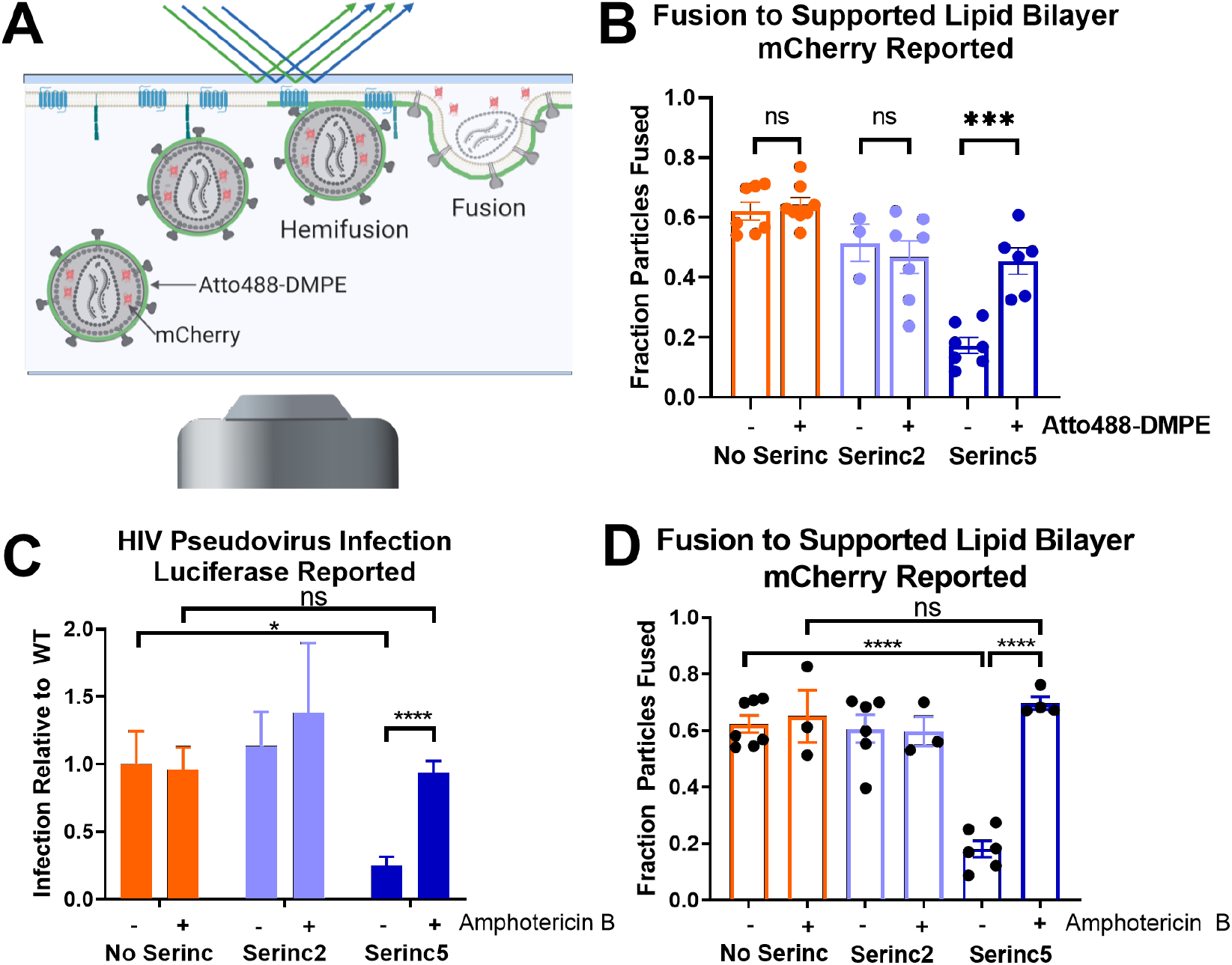
Lipid composition of the viral membrane affects Serinc restriction of HIV membrane fusion. (**A**) Diagram of TIRF microscopy-based single HIV pseudovirus particle fusion assay. Pseudovirus binding to a supported planar plasma membrane containing CD4 receptor and CCR5 co-receptor is reported as a sudden appearance of bright puncta that remain stationary for several imaging frames. Fusion is reported by a decrease in fluorescence over several imaging frames as a genetically encoded content marker, mCherry, diffuses away. (**B**) Incorporation of Atto488-DMPE increases fusion of HIV pseudovirus particles containing Serinc5 but does not affect fusion of particles without Serinc or containing Serinc2. Each data point represents the fraction of particles fused on a separately prepared bilayer. Pre-treatment with 1 μM Amphotericin B increases (**C**) infection and (**D**) fusion of Serinc5 containing HIV pseudoviruses. Data reproduced from (Ward et al., 2020), with permission. *, *p* < 0.05; **, *p* < 0.01; ****, *p* < 0.0001; *ns*, not significant by multiple unpaired t-tests via the Holm-Sidak method. Comparisons not shown are not significant. Each condition includes data from at least three distinct preparations of pseudovirus.

Additionally, we showed that treatment with the polyene antifungal, Amphotericin B, which has complex interactions with membranes, increases infection and fusion of Serinc5-containing HIV pseudoviruses (Fig 3C and D, reproduced from Ward et al., 2020). Similarly, labeling HIV pseudoviruses with the fluorescent lipid, Atto488-dimyristoylphosphatidylethanolamine (DMPE), increased fusion capability of Serinc5-containing HIV pseudoviruses (Fig. 3B). DMPE modified at the headgroup with Atto488 is chemically distinct from unmodified DMPE, so we assessed the effects of exogenous unmodified DMPE on fusion of HIV pseudoviruses incorporating Serinc5, Serinc2, or no Serinc (Fig. 4A) and found the same reversal of Serinc5’s inhibition of HIV membrane fusion. Additionally, we compared the effects of acyl chain length and saturation by examining the fusion of pseudoviruses enriched with dipalmitoyl (DP), dioleoyl (DO), and palmitoyl-oleoyl (PO) PE. Both saturated (DMPE and DPPE) and unsaturated (DOPE) lipids increased fusion of Serinc5-containing pseudoviruses with only minimal effect on pseudoviruses without Serincs or with Serinc2. Thus, restoration of fusion of Serinc5-containing pseudoviruses upon enrichment with PE is a function of the PE headgroup independent of acyl chain length or saturation.

**Fig. 4.**
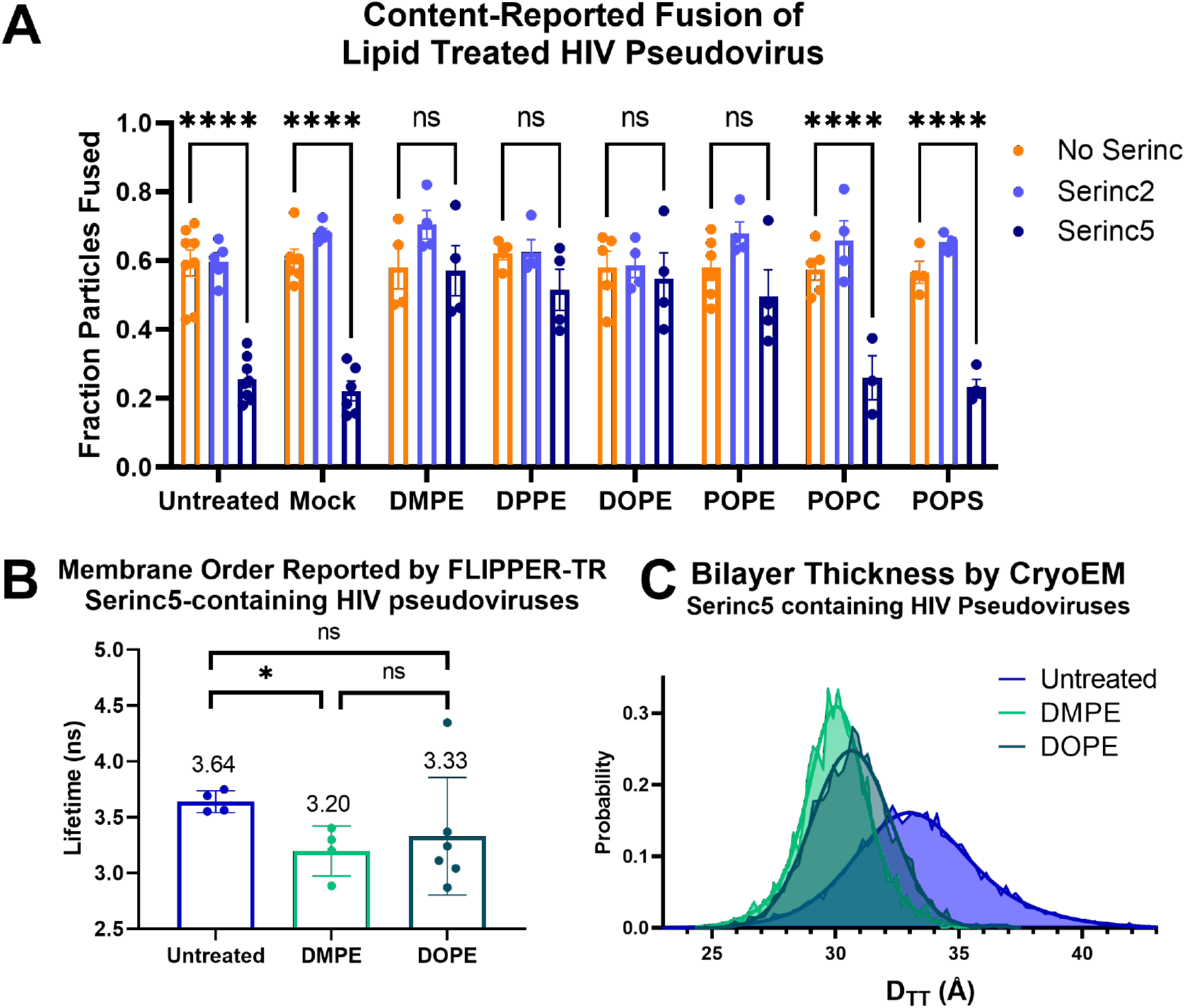
Enrichment of HIV pseudoviruses with PE overcomes restriction of fusion of Serinc5-containing particles by decreasing membrane heterogeneity and order. HIV pseudoviruses with or without Serincs were enriched with the lipids listed on the X-axis, separated from free lipids, and (**A**) fusion with supported plasma membranes was assessed by TIRF microscopy-based single-particle membrane fusion. A two-way ANOVA test was conducted to examine the statistical significance of the effects of Serinc incorporation and lipid enrichment on HIV pseudovirus fusion. There was a statistically significant interaction between the two factors, F(14, 88)=4.894, p<0.001. Tukey’s multiple comparisons test showed very significant differences in the means of the comparisons shown above (**** p<0.0001, ns not significant). None of the No Serinc-Serinc2 comparisons were significant. Each data point represents the fraction of particles fused on a separately prepared supported membrane. Each condition includes data from at least three distinct preparations of pseudovirus. (**B**), Fluorescence lifetimes of FLIPPER-TR in lipid-enriched, Serinc5-containing HIV pseudoviruses. HIV pseudoviruses containing Serinc5 were stained with FLIPPER-TR after enrichment with the lipid listed on the X-axis and imaged by fluorescence microscopy as in Figure 1. Each data point is the result obtained from one field of view within a sample and is plotted with mean and standard deviation. Statistical significance by Welch’s unpaired t-test: * p<0.05, ns not significant. (**C**) Normalized distributions of membrane thickness (DTT) measurements of 5 nm segments of the same preparation of Serinc5-containing HIV pseudoviruses enriched with the lipid listed in the key as panel (**B**) and their corresponding double-Gaussian fits. The unenriched Serinc5 distribution is replotted from Figure 2E for comparison.

To further confirm that the PE headgroup is critical to counteract the fusion restriction of Serinc5, we enriched particles with palmitoyl-oleoyl-phosphatidylcholine (POPC) or palmitoyl-oleoyl-phosphatidylserine (POPS) and found that these lipids had no effect on fusion of any pseudoviruses tested (Fig. 4A). While fewer particles incorporated fluorescently labeled PC than PE or PS (Fig. S6A and C) and the amount of PC incorporated was lower than the amount of PE or PS (Fig. S6B), PS incorporation was equal to PE incorporation yet still failed to increase fusion of Serinc5-containing particles, indicating the specificity of PE in overcoming Serinc5-restriction of HIV membrane fusion. Due to their smaller headgroup, negative membrane curvature-inducing and hydrogen bonding capability, PEs are known to alter the lateral pressure profile and hence membrane bending elasticity of lipid bilayers (Fan et al., 2016).

To better understand the mechanisms underlying the reversal of Serinc5’s inhibition of HIV fusion by PE enrichment, we applied the FLIM and cryoEM assays described above to assess membrane order of Serinc5-containing pseudoviruses that had been enriched with PEs of varying chain length and saturation (Fig. 4B-C and Fig. S7). DMPE is a lipid with two saturated 14-carbon chains and DOPE is an unsaturated lipid with two 18-carbon chains with one *cis* double bond each. Compared to unenriched Serinc5-containing pseudovirus particles, DMPE and DOPE-enriched particles had a clear trend to shorter FLIPPER-TR lifetimes with enrichment with DMPE showing a significant difference from unenriched (Fig. 4B). Imaging the same PE-enriched viral particles by cryoEM revealed there was a marked decrease in membrane thickness (as measured by DTT) for both PEs (Fig. 4C), echoing the decrease in membrane order reported by FLIPPER-TR. Additionally, the PE-enriched Serinc5 particles had much lower variance in membrane thickness than unenriched particles (Fig. S7C). From these data, it appears that enrichment with exogenous PE reverses Serinc5’s increase in membrane order and heterogeneity to restore the particles’ membrane fusion capability.

## Discussion

In this study, we sought to better understand the mechanism by which Serincs restrict HIV membrane fusion and hence entry of HIV particles into cells. We previously established that Serincs 3 and 5, but not Serinc2, inhibit fusion pore opening and constrict the widening of the fusion “neck” of virus particles fusing with plasma membranes that were derived from HIV receptor and co-receptor expressing cells (Ward et al., 2020). Changes in the function of Env would be expected to primarily affect the early steps of membrane fusion that are dependent on conformational rearrangements of Env. However, the final step, pore enlargement, is thought to occur after all conformational rearrangements of Env are complete and thus is dependent on membrane properties such as curvature and line tension for energy input (Kozlov and Chernomordik, 2015). Serinc-dependent changes to the properties of the viral membrane could explain the observed defects in fusion pore dilation with viral envelopes containing Serinc 3 and 5.

To reveal more insightful details on the mechanism by which Serincs inhibit virus entry, we employed fluorescence lifetime imaging and cryoEM methods to assess the importance of lipid order in the viral membrane for Serinc5’s inhibition of HIV membrane fusion. While HIV pseudoviral membranes with and without the non-restricting isoform, Serinc2, were nearly identical by FLIPPER-TR-reported membrane order (Fig. 1) and cryoEM-measured membrane thickness (Fig. 2B, 2C, 2D and 2E, top panel), pseudoviral membranes that incorporated Serinc5 were more ordered (Fig. 1) and more heterogeneous (Fig. 2B), and had a broader interparticle thickness distribution by cryoEM (Fig. 2C and 2D), with many more regions of increased and decreased membrane thickness than viral envelopes with Serinc2 or no Serincs (Fig. 2E, middle panel). Three factors could reconcile the broader thickness distributions observed by cryoEM and the increased lipid order measured by fluorescence lifetime imaging. First, as mentioned above, overlapping areas of L_o_ and L_d_ phases that are too small to be resolved in cryoEM projection images will lead to an apparent broadening of the membrane thickness distributions measured by cryoEM. Second, domain sizes may fluctuate in space and time and these fluctuations may be beyond the resolution of cryoEM such that observed thicknesses are blurred over those of fluctuating segments. Third, a more favorable partitioning of FLIPPER-TR into L_o_ than into L_d_ domains or perhaps a more complex behavior of the probe at lipid domain interfaces may explain the increased order measured in the Serinc5 particles. Despite these caveats, we note that Serinc5 increases heterogeneity in the viral membrane with increased probability of both thinner and thicker segments of membrane, causing broadening of the distribution of membrane thicknesses. When this is examined in bulk by FLIM (Fig. 1), the overall effect is an increase in ordered fraction of the membrane and the overall trend of the effect of Serinc5 incorporation as measured by both techniques is skewing of the viral membrane towards increased lipid order. As a change in membrane order only occurred with incorporation of the restricting isoform, Serinc5, and not the non-restricting isoform, Serinc2, this strongly suggests the restriction function of Serinc5 is related to altered lipid order and organization in the viral membrane. Notably, fluorescent membrane order probes that report solvent accessibility, a property frequently but not always related to membrane order (M’Baye et al., 2008), failed to detect increased membrane heterogeneity in Serinc5-containing particles (Raghunath et al., 2022) as we have reported here by two different techniques, one of which is completely label-free,

Our observation that PE increases fusion of Serinc5-containing viruses (Fig. 4A) is further demonstration of the importance of altered membrane properties for Serinc5-mediated fusion inhibition. The reversal of Serinc5’s function with PE enrichment is accompanied by a reversal of Serinc5’s effects on viral membrane organization with decreased order reported by FLIPPER-TR (Fig. 4B) and dramatically reduced membrane thickness by cryoEM (Fig. 4C). Again, these two techniques report on slightly different properties of ordered/disordered membranes, so while the magnitude of the effect of PE enrichment is different between the two measures, the overall direction of the change is consistent—PE enrichment reduces membrane order regardless of acyl chain length or saturation. The data strongly suggest that altered membrane order and heterogeneity is essential for Serinc5’s restriction of HIV membrane fusion.

The observation of coexistence of thicker and thinner membrane domains in HIV pseudoviral membranes (Fig. 2 and Fig. S3) is, to our knowledge, the first time that domain coexistence has been directly observed in a viral membrane. Previous bulk measurements of lipid order of HIV particles have shown the membrane to be largely ordered, but that lipid order in HIV particles also depends on the producing cell line (Lorizate et al., 2009). Phase separation has been observed in supported lipid bilayers made from viral membrane lipid extracts (Huarte et al., 2016), but not in intact viral particles. Building on our observation of membrane heterogeneity in the HIV envelope, we note a line of density (Fig. S4) at a distance from the bilayer consistent with an incomplete MA layer of HIV (Eells et al., 2017; Qu et al., 2021). Prior reconstitution experiments have shown thinning of the bilayer upon MA binding to model membranes (O’Neil et al., 2016) and similarly, we observe MA density most frequently under thinner portions of the viral membrane, thus recapitulating prior reconstitution data *in virio.* It is possible that the underlying MA layer introduces tension and further changes the mechanical properties of the viral envelope. However, a more detailed analysis of MA dependent effects on the viral envelope is beyond the scope of the current study.

### Ideas and Speculation

It has been shown previously that both Env and Serinc5 partition into membrane domains with more ordered lipids (Schulte et al., 2018; Schwarzer et al., 2014) and that Serinc5 disrupts the clustering of Env in the viral membrane (Chen et al., 2020). If clustering in the mature viral envelope is driven by confinement of Env to small, ordered lipid domains, increasing area or fragmentation of ordered domains could reduce clustering, thus decreasing the number of Env trimers available to catalyze fusion (Brandenberg et al., 2015). Additionally, Serincs have been hypothesized to alter Env conformation as demonstrated by multiple methods (Beitari et al., 2017; Chen et al., 2020; Kirschman et al., 2022; Leonhardt et al., 2022; Schulte et al., 2018; Sood et al., 2017; Staropoli et al., 2019). Changes in Env conformation, especially increased exposure of the membrane proximal external region (MPER) of Env, could also be a result of altered membrane thickness and order (Hollingsworth et al., 2018).

As a result of increased order, the Serinc5 viral membrane is likely stiffer than a viral membrane without Serincs (Steinkühler et al., 2019), potentially explaining arrest of membrane fusion at highly curved intermediates (Fig. 5). Additionally, changing area fraction and distribution of ordered lipid domains may reflect altered line tension between domains in the viral membrane (Usery et al., 2017). Previous work from our laboratory has shown that in phase separated viral and target membranes, the energy of line tension can drive HIV gp41-mediated fusion, that this effect depends on the size and distribution of ordered domains in the target membrane, and that increasing ordered domain area in a viral particle beyond its optimum would be expected to inhibit fusion (Yang et al., 2016a). Applying this simplified model to study the effect of increased ordered domain area in a membrane of viral particles with 110 nm diameter, we found that increasing the fraction of the membrane in an ordered domain first increases and then sharply decreases the contribution of line tension to the free energy change of fusion (Fig. S8). It is well known that the final step of fusion, fusion pore expansion, is dependent on lateral tension in the membrane (Kliesch et al., 2017; Kozlov and Chernomordik, 2015; Ryham et al., 2016; Shillcock and Lipowsky, 2005; Staykova et al., 2011). In a more ordered membrane, the elastic modulus would be expected to change along with altered curvature and line tension, all of which affect the energy profile for membrane fusion (Fan et al., 2016; Siegel, 2008). Decreased lateral tension would also be expected to inhibit fusion pore expansion resulting in the increased frequency of early fusion products observed with cinched membranes (Ward et al., 2020).

**Fig. 5.**
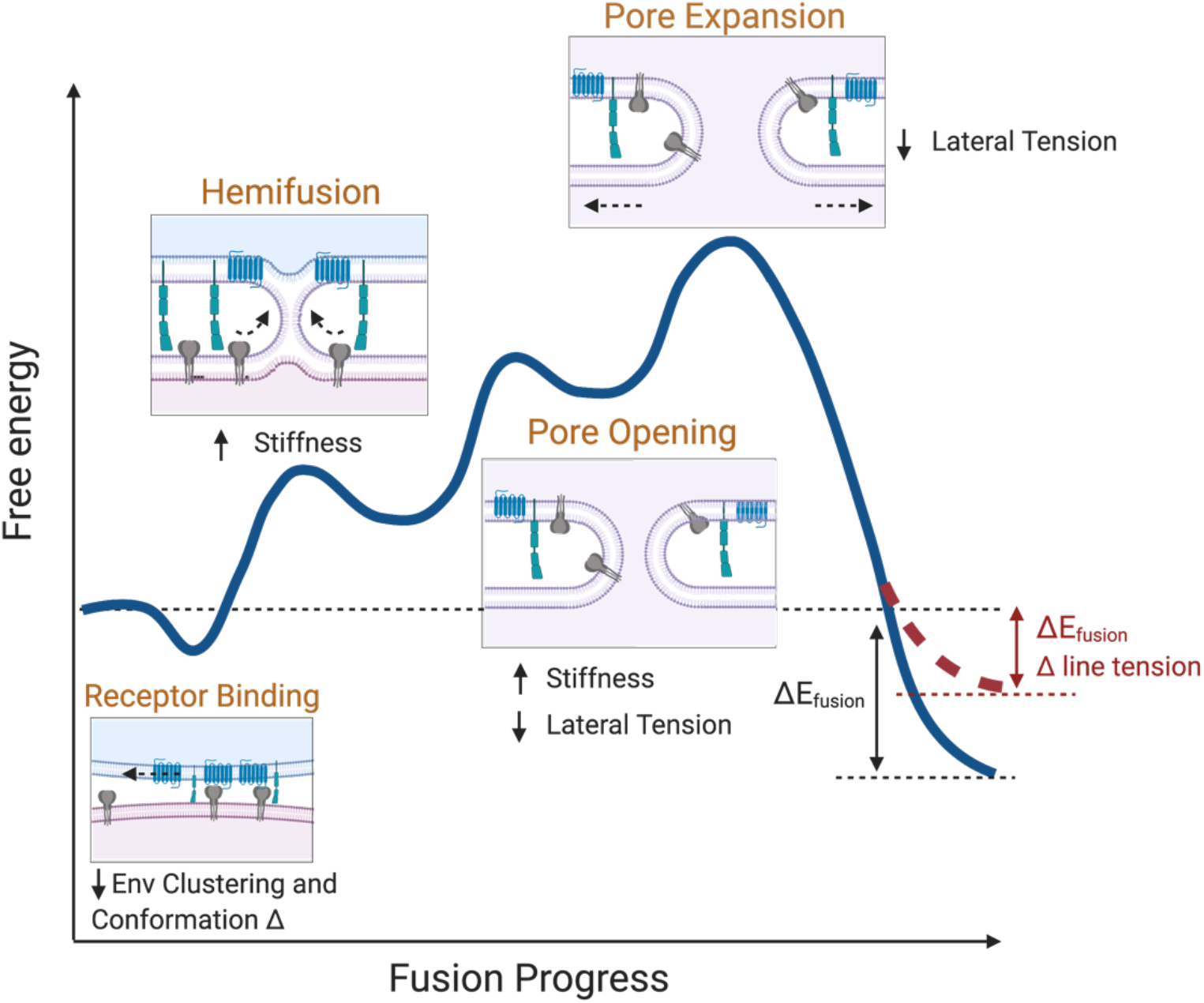
Model of energy changes to fusion resulting from Serinc5-induced increase in membrane heterogeneity of the viral envelope. The free energy change of intermediate steps (orange titles) along the fusion reaction is shown by the blue line. Proposed effects of Serinc5-induced alterations of membrane physical properties are listed in black near each intermediate state. Many of these properties are inter-related and are not easily disentangled. The effect of increased membrane heterogeneity on the contribution of line tension to the energetics of fusion is shown by the red dashed line.

Serinc 3 and 5’s possible functions as lipid binding and/or translocating integral membrane proteins (Leonhardt et al., 2022; Pye et al., 2020; Trautz et al., 2017) could alter effective lipid concentrations and asymmetry in the viral membrane, and thus alter lipid order. Specifically, Serincs 3 and 5 may alter the distribution of PE across the two leaflets of the viral membrane. PE is known to enhance membrane fusion, especially when preferentially incorporated into the two contacting leaflets of the two fusing membranes (Chernomordik and Kozlov, 2005; Churchward et al., 2008; Kreutzberger et al., 2017). As a cone-shaped molecule, PE induces frustrated negative curvature in the pre-fusion membrane, increasing lateral tension in the headgroup region (Fan et al., 2016). In support of this hypothesis, we showed repletion of PE in the viral outer leaflet can overcome Serinc5’s inhibitory function on fusion (Fig. 4A) and that this was accompanied by larger changes in membrane order that likely promote fusion as well. The fundamental and unifying change underlying the potential hypotheses for Serinc 3 and 5’s mechanism of action is increased viral membrane heterogeneity.

Previously, we hypothesized that Serinc5 would need to cause “broad energetic changes” to the membrane fusion process to accumulate arrested intermediates at every step, as we observed (Ward et al., 2020). With the data presented here, we show that Serinc5 increases viral membrane heterogeneity and that this is necessary for the partial inhibition of HIV fusion. Contextualizing previous observations and hypotheses about Serinc5’s mechanism of action with our data, we posit that alteration of membrane order underlies the other effects of Serinc5 incorporation into the HIV envelope. Following from this, it bears further study whether agents that increase membrane order could be effective anti-HIV therapeutics. With the recent discovery that Serinc5 restricts SARS-CoV-2 membrane fusion (Timilsina et al., 2022), it is possible that therapeutics based on a similar mechanism could also be effective against SARS-CoV-2 infection.

## Materials and Methods

### Cell lines, reagents, and plasmids

HEK 293T/17 cells (ATCC) were maintained in high glucose Dulbecco’s Minimum Essential Media (Gibco) supplemented with 10% fetal bovine serum (Atlanta Biologicals), 1% antibiotic-antimycotic (Gibco), 1 mM sodium pyruvate (Gibco), and 2 mM glutamine (Gibco). CD4 and CCR5 overexpressing HeLa cells (gift of David M. Rekosh, University of Virginia) were maintained in Iscove’s Modified Dulbecco’s Medium (Gibco) supplemented with 10% fetal bovine serum and 1% antibiotic/antimycotic with 0.5 mg/mL of G418 (Gibco), and 1 µg/mL puromycin. All cells were maintained at 37°C with 5% CO_2_ atmosphere.

pHIV-luciferase, pHIV-Rev, and pHIV-pack were gifts of Wen Yuan (University of Virginia). pHIV-Env-SF162 was provided by the AIDS Reagent Program. pHIV-imCherry (Padilla-Parra et al., 2013) was a gift of Gregory Melikian (Emory University). pPBJ5-Serinc2-HA was a gift of Massimo Pizzato (University of Trento) and pPBJ5-Serinc5-HA was a gift of Heinrich Gottlinger (University of Massachusetts Medical School, Worcester).

The following compounds were purchased from Avanti Polar Lipids and used without modification: brain phosphatidylcholine (bPC), egg sphingomyelin (SM), 1-palmitoyl-2-oleoyl-sn-glycero-3-phosphoethanolamine (POPE), 1,2-dioleoyl-sn-glycero-3-phosphoethanolamine (DOPE), 1,2-dimyristoyl-sn-glycero-3-phosphoethanolamine (DMPE), 1,2-dipalmitoyl-sn-glycero-3-phosphoethanolamine (DPPE), 1-palmitoyl-2-oleoyl-glycero-3-phosphocholine (POPC), 1,2-dipalmitoyl-sn-glycero-3-phosphocholine (DPPC), 1-palmitoyl-2-oleoyl-sn-glycero-3-phospho-L-serine (POPS), 1-palmitoyl-2-oleoyl-sn-glycero-3-phospho-(1'-rac-glycerol) (POPG), 1-palmitoyl-2-{6-[(7-nitro-2-1,3-benzoxadiazol-4-yl)amino]hexanoyl}-sn-glycero-3-phosphocholine (NBD-PC), 1-palmitoyl-2-{6-[(7-nitro-2-1,3-benzoxadiazol-4-yl)amino]hexanoyl}-sn-glycero-3-phosphoserine (NBD-PS), and 1-palmitoyl-2-{6-[(7-nitro-2-1,3-benzoxadiazol-4-yl)amino]hexanoyl}-sn-glycero-3-phosphoethanolamine (NBD-PE). Cholesterol was purchased from Sigma-Aldrich. 1,2-dimyristoyl-*sn*-glycero-3-phosphoethanolamine-PEG3400-triethoxysilane (DPS) was synthesized as described previously (Wagner and Tamm, 2000).

### HIV pseudovirus preparation

HIV pseudoviruses were produced as described before by transfection of HEK 293T cells (ATCC) with Lipofectamine 2000 (Invitrogen) and the following amounts of plasmids per 10 cm dish: 13 μg pHIV-luciferase, 5 μg pHIV-pack, 4 μg pHIV-Env-SF162 (Cheng-Mayer et al., 1997; Stamatatos et al., 2000, 1998), 4 μg pHIV-imCherry, 1 μg pHIV-Rev and 4μg of pBJ5-Serinc2-HA (Rosa et al., 2015) or pBJ5-Serinc5-HA (Usami et al., 2015) as indicated in text. Culture media was changed 4-6 hours after transfection to phenol-red free DMEM supplemented with 10% FBS, 1% antibiotic-antimycotic, 1 mM sodium pyruvate, and 2 mM glutamine. Culture supernatants were harvested 2 days after transfection and cleared by centrifuging 5000xg before passing through a 0.22 μm filter. HIV pseudoviruses were pelleted through a 25% sucrose-HME (20 mM HEPES, 20 mM morpholineethanesulfonic acid [MES], 130 mM NaCl, 1 mM EDTA [pH 7.4]) cushion as previously described (Hulseberg et al., 2019) and resuspended in buffer HME without sucrose. Pseudovirus preparations for cryoEM were further purified by density dependent centrifugation on a discontinuous sucrose gradient composed of 65% sucrose-HME and 25% sucrose-HME spun 151,000xg for 18 hours. Pseudovirus was collected from the 65%/25% sucrose interface, diluted in buffer HME without sucrose, and repelleted through a 25% sucrose cushion. After resuspension in buffer HME without sucrose, the pseudovirus preparation was aliquoted and stored at −80°C. Additionally, the concentration of HIV p24 in each preparation was measured by ELISA (Toohey et al., 1995; Wehrly and Chesebro, 1997) and used to normalize the amount of pseudovirus added to downstream experiments. Quantification of Serinc incorporation into pseudoviral particles prepared in the same manner is shown in supplemental figures 3 and 8 of (Ward et al., 2020). While Serinc5 incorporates into viral particles more readily than other Serincs, there is no relationship between level of Serinc incorporation and restriction activity (Diehl et al., 2019) for Serinc levels above a low threshold (Ward et al., 2020).

### Lipid enrichment of viruses

10^−10^ moles of the indicated lipid were dried on the bottom of a glass test tube to remove chloroform/methanol solvent and resuspended by vigorous vortexing in buffer HB (20 mM HEPES, 150 mM NaCl [pH 7.4]) to yield a concentration of 1.4 μM. 21 ng of HIV pseudovirus, as measured by p24 ELISA, was mixed with the lipid suspension. The mixture was incubated at room temperature for 2 hours on a rotary spinner. To remove free lipid, the HIV pseudovirus mixture was diluted up to 1.5 mL in buffer HB and pelleted by spinning at 21,000xg for 1 hour at 4°C before final resuspension in buffer HB. Lipid-enriched HIV pseudoviruses were used within 24 hours.

### Large unilamellar vesicle preparation

Chloroform stocks of desired lipids were mixed and the solvent was evaporated under a gentle stream of nitrogen gas. The resulting lipid film was desiccated under vacuum for at least one hour before resuspension in buffer appropriate to the experiment to a final concentration of 1 mM. After vortexing at room temperature, the lipid suspension was subjected to 10 freeze/thaw cycles in liquid nitrogen and warm water before extrusion through two 100 nm polycarbonate membranes (Avestin). Resulting LUVs were stored at 4°C and used within 24 hours of extrusion.

### Plasma membrane bleb preparation

Blebs were produced from HeLa cells overexpressing CD4 and CCR5 by previously published methods (Sezgin et al., 2012; Yang et al., 2017). Briefly, when cells reached 90% confluence, they were washed twice with blebbing buffer (10 mM HEPES, 150 mM NaCl, 2 mM CaCl_2_, pH 7.4) and blebbing was induced by replacing buffer on the cells with 5 mL of 25 mM formaldehyde (J.T.Baker) and 2 mM dithiothreitol (DTT) diluted in blebbing buffer and incubating the cells at 37°C, 5% CO_2_ for 1 hour. After an hour, blebs were detached from cells by shaking on a radial shaker at room temperature for 1 hour before the supernatant was collected and cleared of large cell debris by centrifuging at 100xg for 10 minutes. Blebs were pelleted at 20,000xg for 1 hour and washed twice in blebbing buffer without DTT or formaldehyde.

### TIRF supported lipid bilayer fusion assay

Supported planar plasma membranes derived from blebs were prepared as previously described (Kalb et al., 1992; Wagner and Tamm, 2000; Yang et al., 2017). Quartz slides were cleaned in piranha solution (95% H_2_SO_4_ and 30% H_2_O_2_ in a 3:1 ratio) and rinsed in 12 liters of deionized water. Next, a lipid monolayer composed of 4:1 brain phosphatidylcholine and cholesterol (Avanti Polar Lipids) with 3% 1,2-dimyristoyl-*sn*-glycero-3-phosphoethanolamine-PEG3400-triethoxysilane was deposited on the quartz slide by the Langmuir-Blodgett method. A chloroform solution of the lipid mixture was applied to a Nima 611 Langmuir-Blodgett trough and after letting the solvent evaporate for 10 minutes, the lipid layer was compressed at a rate of 10 cm^2^/min to a pressure of 32 mN/m. A cleaned, rinsed, and dried quartz slide was rapidly dipped (68 mm/min) and slowly removed (5 mm/min) from the trough and then dried in a desiccator chamber overnight.

The slide was then assembled into a custom-built microscopy flow cell and plasma membrane blebs diluted in blebbing buffer without DTT or formaldehyde were flowed in to form the outer leaflet of the supported planar plasma membrane. After 1-2 hours at room temperature, the flow cell was washed with multiple volumes of blebbing buffer, then multiple volumes of buffer HB, and transferred to a prism-based TIRF microscope (Zeiss AxioObserver Z1). The sample was excited with a 561nm diode laser (OBIS 561 nm LS, Coherent) at an angle of 72 degrees from normal and emission light was filtered through a dichroic mirror (DC565, Semrock) and a band-pass filter (BP605/50, Semrock). Video was recorded by an EMCCD (DV887ESC-BV, Andor Technology) in frame transfer mode with an exposure time of 0.2 s for 13.3 minutes as a dilution of HIV pseudovirus totaling 21 ng of p24 as measured by ELISA was flowed into the chamber. Laser intensity, shutter, and camera were controlled by a custom LabView program (National Instruments).

Intensities of single particles over time were extracted with a custom-built LabView program and classified as representing binding without fusion or binding with fusion based on the criteria described in (Ward et al., 2020).

### FLIM data acquisition and analysis

HIV pseudoviruses and LUVs were stained with 500 nM FLIPPER-TR (Cytoskeleton Inc.) for 2 hours at room temperature. Pseudoviruses and LUVs were resuspended in buffer HB and added to a poly-L-lysine coated coverslip. After liposomes and pseudoviruses were allowed to settle and adhere to the coverslip for 30 minutes, unbound sample was washed in buffer HB. Samples were imaged on a Leica Stellaris8 microscope with an 80 MHz pulsed white light laser, HyD detector, and FALCON software. Background was subtracted by intensity thresholding and the fluorescence lifetime of all pixels above the intensity threshold within the field of view was fit with two components per the n-component reconvolution with IRF function within Leica LAS-X software. Only the longer component is displayed as this is the more sensitive reporter of membrane order and tension (Colom et al., 2018). For data collected in bulk (Fig. S2), LUVs were stained in the same manner but added to a quartz cuvette instead of a coverslip and spectra were acquired in a Fluorolog-QM 75-22-C (Horiba, Canada) spectrofluorometer with a SuperK Extreme high power super continuum white laser (NKT Photonics) in time-correlated single photon counting mode. Laser repetition rate was 5.5 MHz. Spectra were again fit to a two-component exponential curve with background subtraction of a buffer only spectrum with the integrated PowerFit-10 decay analysis package (Horiba).

### CryoEM of HIV pseudovirus particles

C-Flat 2/2-3C or 1.2/1.3-3C grids (Electron Microscopy Sciences) were glow discharged at 10 mA for 90 seconds. A suspension of HIV pseudovirus in buffer HME was applied and blotted from the grid before freezing in liquid nitrogen cooled ethane. For initial experiments, LUVs composed of 20/40/35/5 Chol/DPPC/DOPC/POPG were spiked into the pseudovirus suspension before freezing to serve as internal standards for quality control. The grids were imaged on a Titan Krios electron microscope operating at 300 kV equipped with a K3/GIF (Gatan) and controlled by EPU software (ThermoFisher Scientific). Magnification was 33,000X, which yielded a pixel size of 2.7 Å. As described in (Heberle et al., 2020), the optimal total dose was set at 13.8 e/Å^2^. Micrographs were motion corrected with MotionCorr2 (Zheng et al., 2017) (10 by 10 patch for 10 iterations with a tolerance of 0.5, dose weighted) before analysis of trough-to-trough distance as described previously (Heberle et al., 2020).

### Calculation of the contribution of line tension to free energy change of membrane fusion

The previously described model to calculate the free energy change due to changes in line tension during fusion between phase separated membranes (Yang et al., 2016a) was adapted to the parameters relevant to the presented experimental data. Specifically, fusion between a flat membrane and a vesicle of 110 nm diameter with one L_o_ and one L_d_ domain each, representing the plasma membrane and HIV viral membrane respectively. Assuming simple geometries without domain fluctuations, the boundary energy of an isolated domain is given by E = γL, where γ is the line tension and L is the circumference of the domain (Kuzmin et al., 2005). Line tension in both membranes is assumed to be 1 pN (Kuzmin et al., 2005) and the negative energy gain from line tension reduction (dE) relative to *k_B_T* (-dE/ *k_B_T*) was calculated for varying lipid domain sizes and fractions in both membranes. Depending on vesicle size, area fraction of L_o_ phase membrane, and domain sizes in the target membrane, fusion of the two membranes can result in an energy gain or energy cost from changes in line tension (Fig. S8).

### Statistical Analysis

All tests of statistical significance calculated in GraphPad Prism 9. Specific tests are listed in the figure legends.

## Acknowledgments

We thank Drs. Barbie Ganser-Pornillos and Kelly Dryden for expert help with the collection of the cryo-EM data acquired at the Molecular Electron Microscopy Core at the University of Virginia, which is supported in part by NIH grant U24 GM116790. Additionally, we thank the lab of Dr. Cliff Stains for assistance with the collection of FLIM spectra.

## Funding

National Institutes of Health grant F30 HD101348 (AEW)

National Institutes of Health grant R01 AI30557 (LKT)

National Institutes of Health grant R01 GM138887 (FAH and MNW)

National Institutes of Health grant R35 GM134949 (IL)

National Institutes of Health grant R01 GM120351 (KRL)

National Institutes of Health grant T32 GM080186 (AEW)

National Institutes of Health grant T32 GM007267 (AEW)

Volkswagen Foundation Grant 93091 (IL)

William Wheless III professorship (MNW)

## Author contributions

Conceptualization: AEW, MNW, FAH, IL, JMW, VK, LKT

Methodology: AEW, MNW, FAH, IL, KL, VK, LKT

Investigation: AEW, DS, MNW, FAH, VK, IL, KL

Visualization: AEW, FAH, MNW, VK

Supervision: JMW, LKT

Writing—original draft: AEW

Writing—review & editing: all authors

## Competing interests

Authors declare that they have no competing interests.

## Data and materials availability

All data, code, and materials used in the analyses available upon request

## Supplementary Materials

**Fig. S1.**
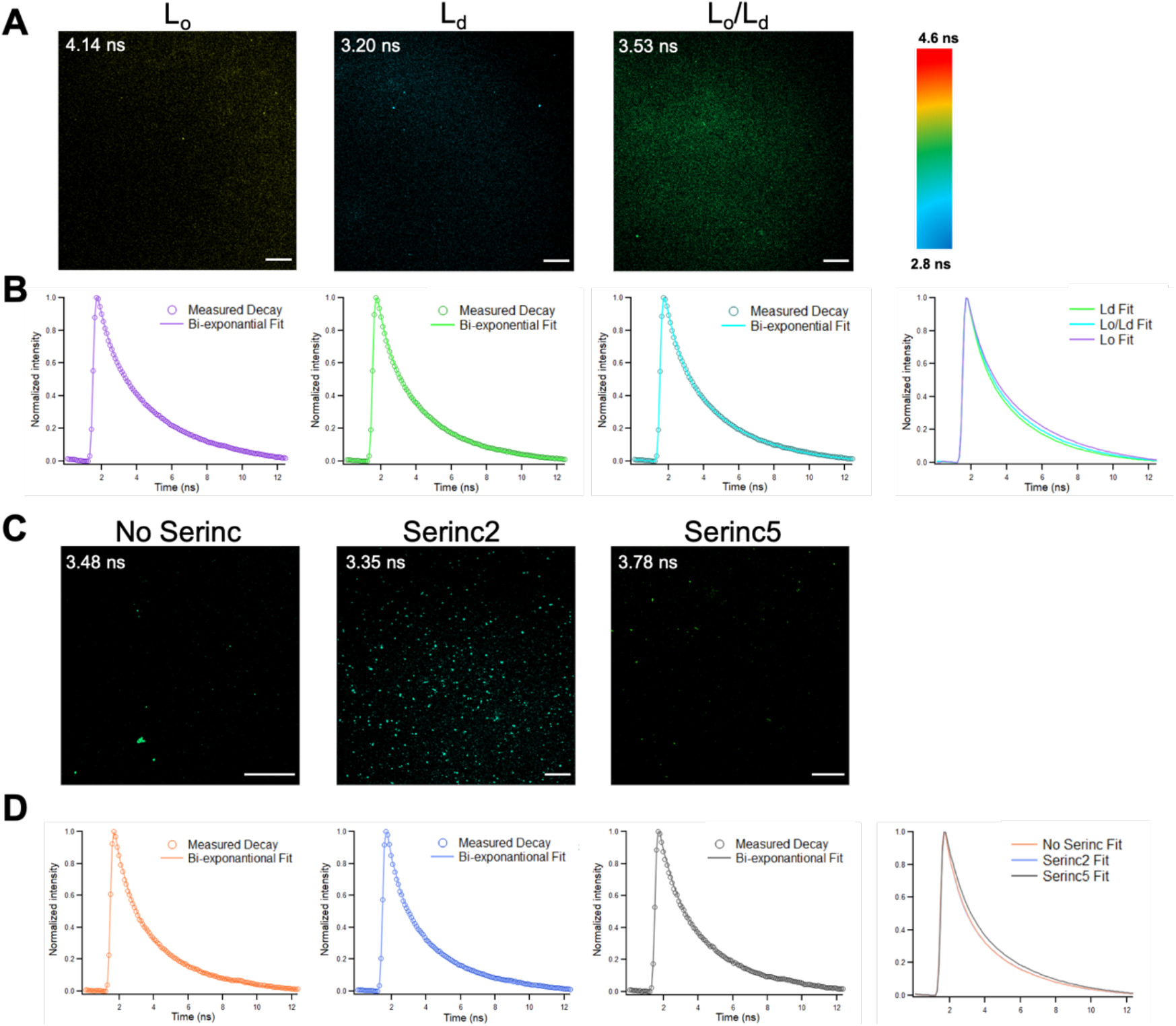
Fluorescence Lifetime Microscopy of FLIPPER-TR stained LUVs and HIV pseudoviruses. (**A**) Example micrographs of large unilamellar liposomes (LUVs) of compositions known to exist solely in L_o_ (35/40/25, SM/chol/POPC), solely L_d_ (25/5/70, SM/chol/POPC), or as a mixture of L_o_ and L_d_ (40/20/40, SM/chol/POPC) phases that were stained with FLIPPER-TR and adhered to a poly-L-lysine coverslip. Lifetime is scaled in color as shown in the key to the right. (**B**) Decay curves of all pixels in the micrograph above that exceed an intensity threshold to exclude background but include punctate fluorescence of LUVs. LUVs from different samples plotted on same axes in the rightmost plot. (**C**) Example micrographs of HIV pseudoviruses stained with FLIPPER-TR and adhered to a poly-L-lysine coverslip. Lifetime is scaled in color the same as in **A**. (**D**) Decay curves of HIV pseudoviruses calculated as in **B**. The Serinc2 curve is plotted but difficult to visualize as it completely overlaps with No Serinc. Data are examples from same dataset as Figure 1. Scale bars are 20 μm.

**Fig. S2.**
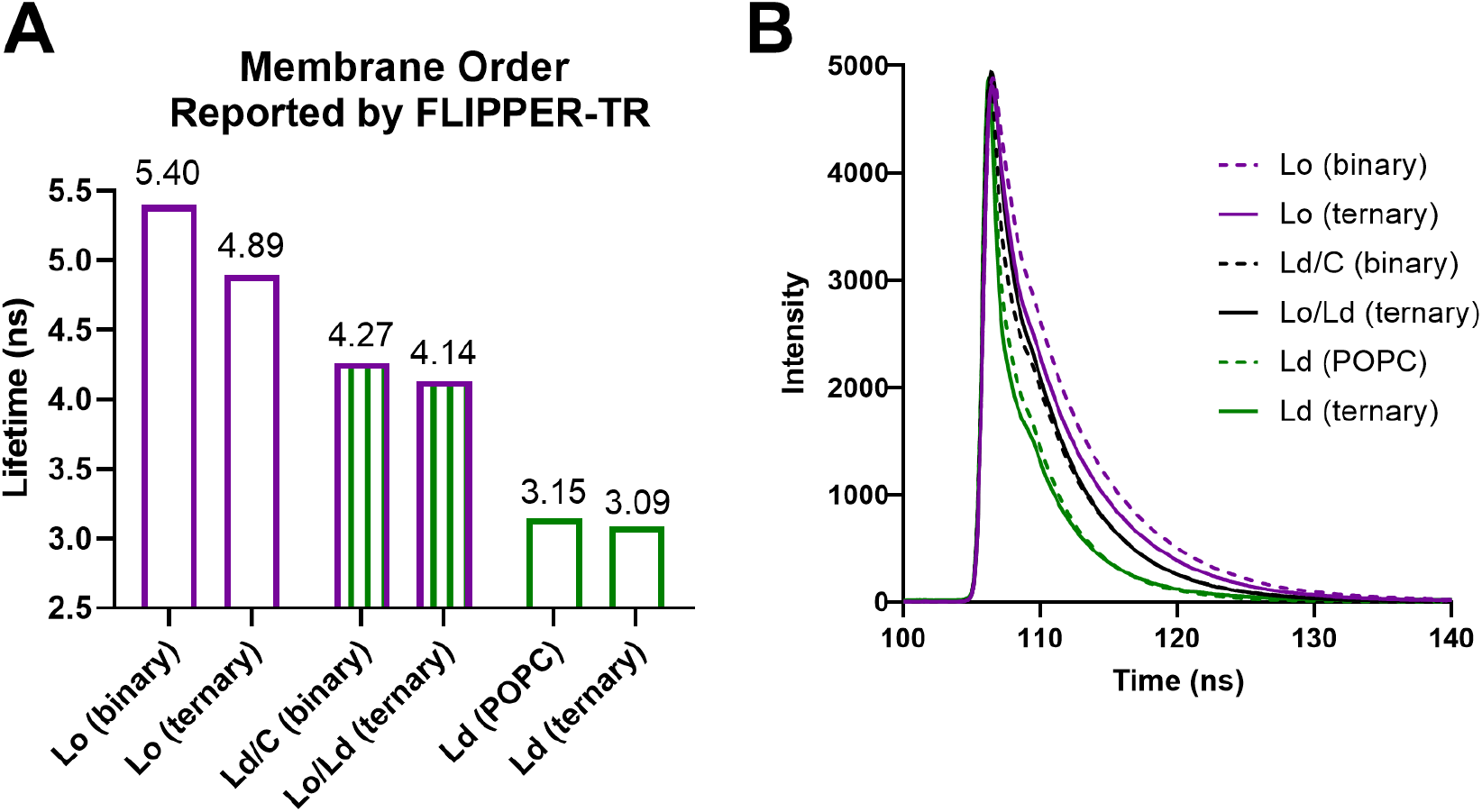
Lifetimes of FLIPPER-TR in LUVs of varying composition. Large unilamellar liposomes (LUVs) of compositions known to exist solely in L_o_ (ternary 35/40/25, SM/chol/POPC; binary 50/50 eggSM/chol), solely L_d_ (ternary 25/5/70, SM/chol/POPC; POPC is solely POPC), L_d_ with cholesterol (L_d_/C binary 40/60 Chol/POPC), or as a mixture of L_o_ and L_d_ (ternary 40/20/40, SM/chol/POPC) phases were stained with FLIPPER-TR as described in materials and methods and lifetimes were measured in a cuvette in a fluorometer (**A**). Decay curves (**B**) were fit to two exponentials, the longer of which is plotted in (**A**) with calculated error (too small to visualize). Data are from a single experiment.

**Fig. S3:**
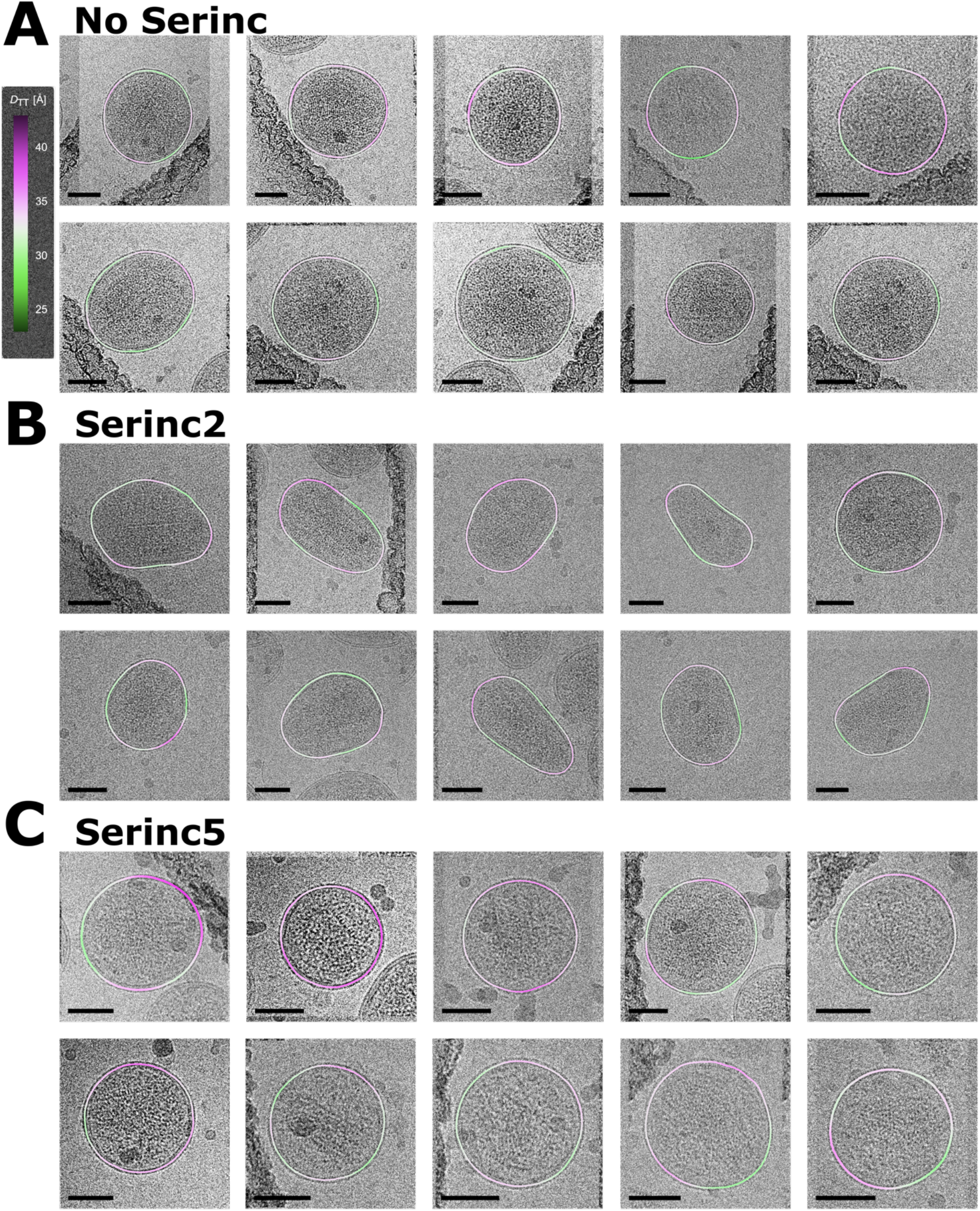
Additional example micrographs with membrane thickness (D_TT_) overlay of HIV pseudovirus particles. Trough to trough distances (D_TT_) of viral membranes are shown as heat maps (scale on the left) centered on the average distance. Micrographs shown from multiple preparations of HIV pseudoviruses without Serincs (**A**), with Serinc2 (**B**), and with Serinc5 (**C**). All black scale bars are 50 nm. The edge of the carbon support film is visible in some micrographs along with small areas of electron dense contaminants. Pixel padding to ensure square images for analysis visible in some micrographs.

**Fig. S4.**
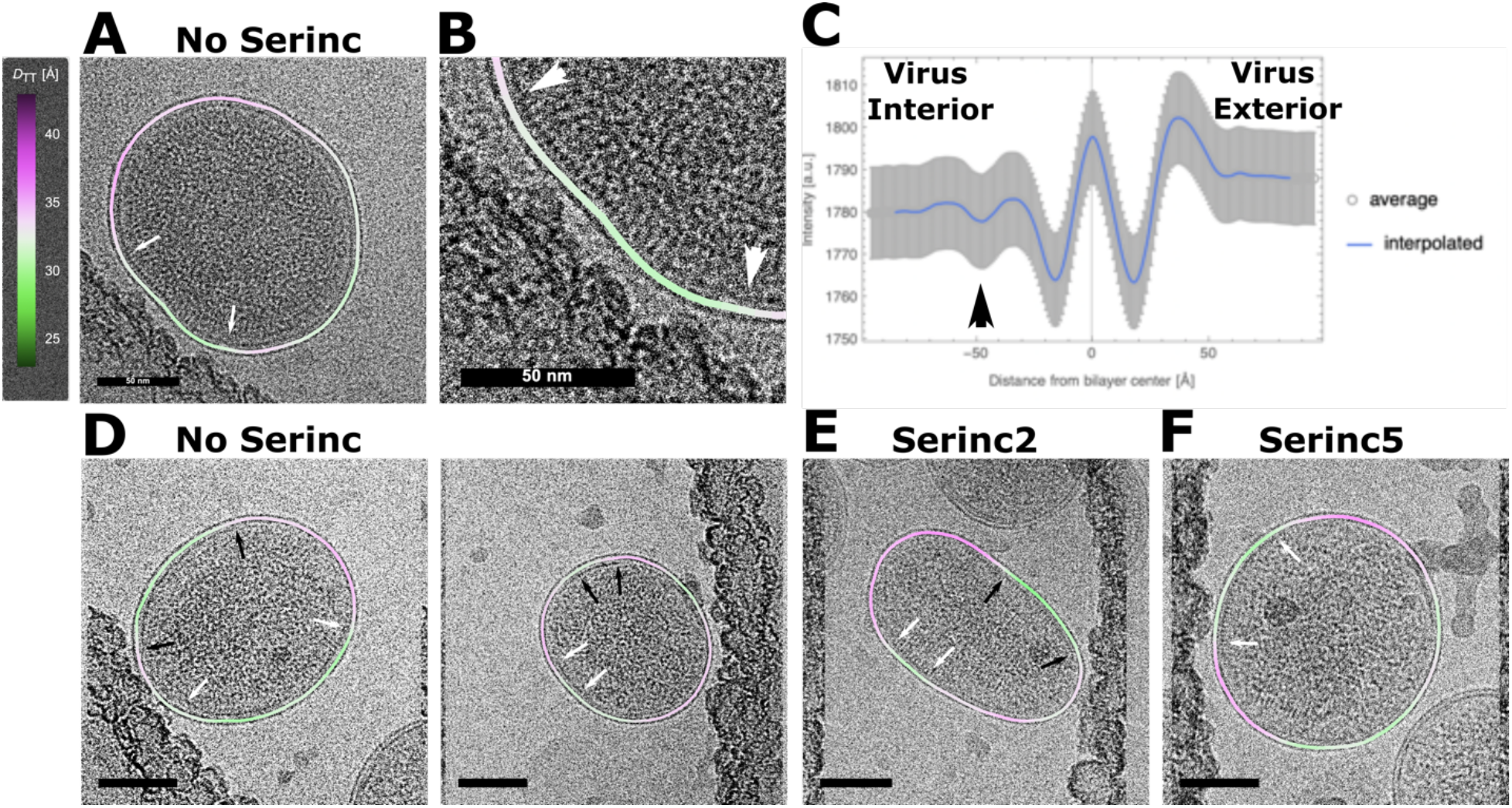
Observation of linear densities preferentially under areas of thinner membranes. Trough to trough distance (D_TT_) of viral membranes shown as heat map (scale on the left) centered on the average distance. Arrows of same color denote edges of discontinuous linear densities. All scale bars are 50 nm. (**A**) Example HIV pseudoviral particle produced without Serincs with close up (**B**) of area between arrows. (**C**) Radially averaged intensity profiles for 104 viral particles without Serincs showing an additional sub-membrane density (black arrowhead) at a distance from the inner membrane leaflet trough consistent with previous measurements (47). (**D-F**) Additional example micrographs of HIV pseudoviral particles without Serincs (**D**), with Serinc2 (**E**), and Serinc5 (**F**) with discontinuous areas of visible membrane-associated density.

**Fig. S5.**
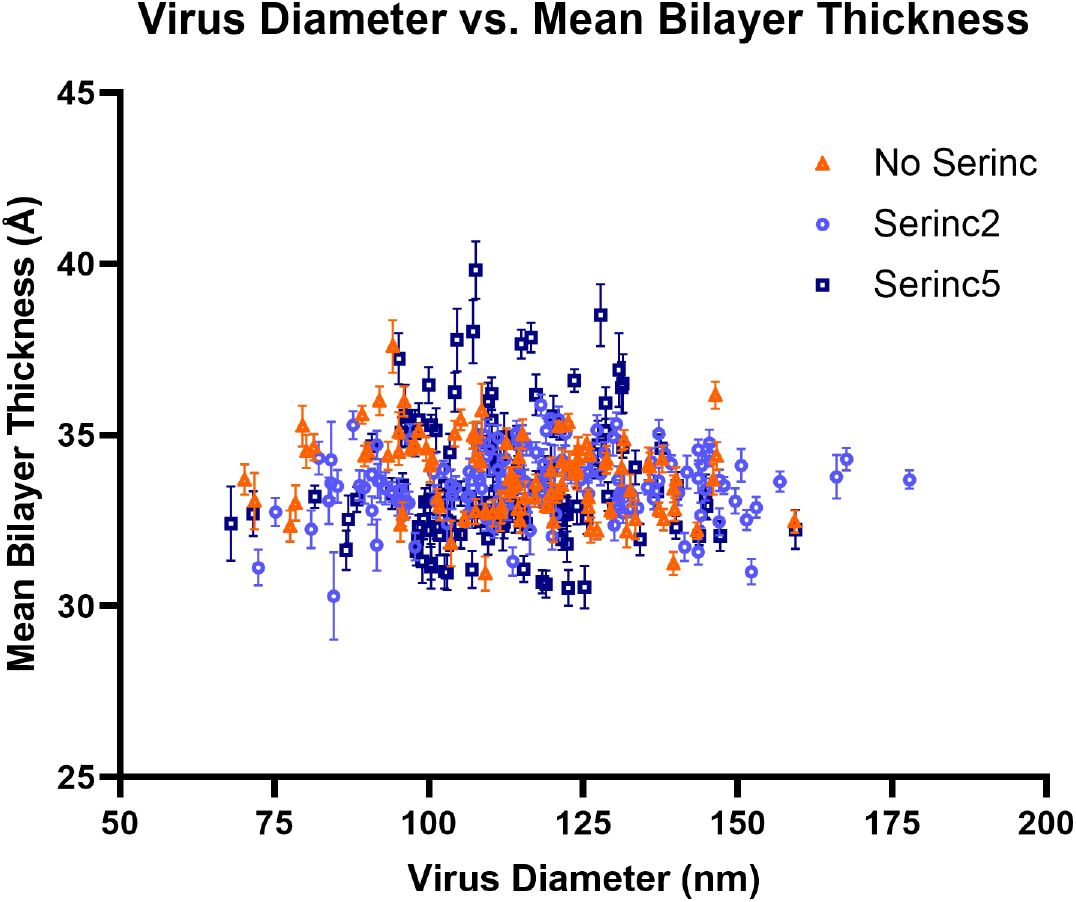
Relationship between size distribution and membrane thickness for HIV pseudoviruses imaged by cryoEM. The diameter of the pseudovirus membrane contour was plotted against the mean bilayer thickness with standard deviation for each particle analyzed in Fig. 2. Each point is a single viral particle.

**Fig. S6.**
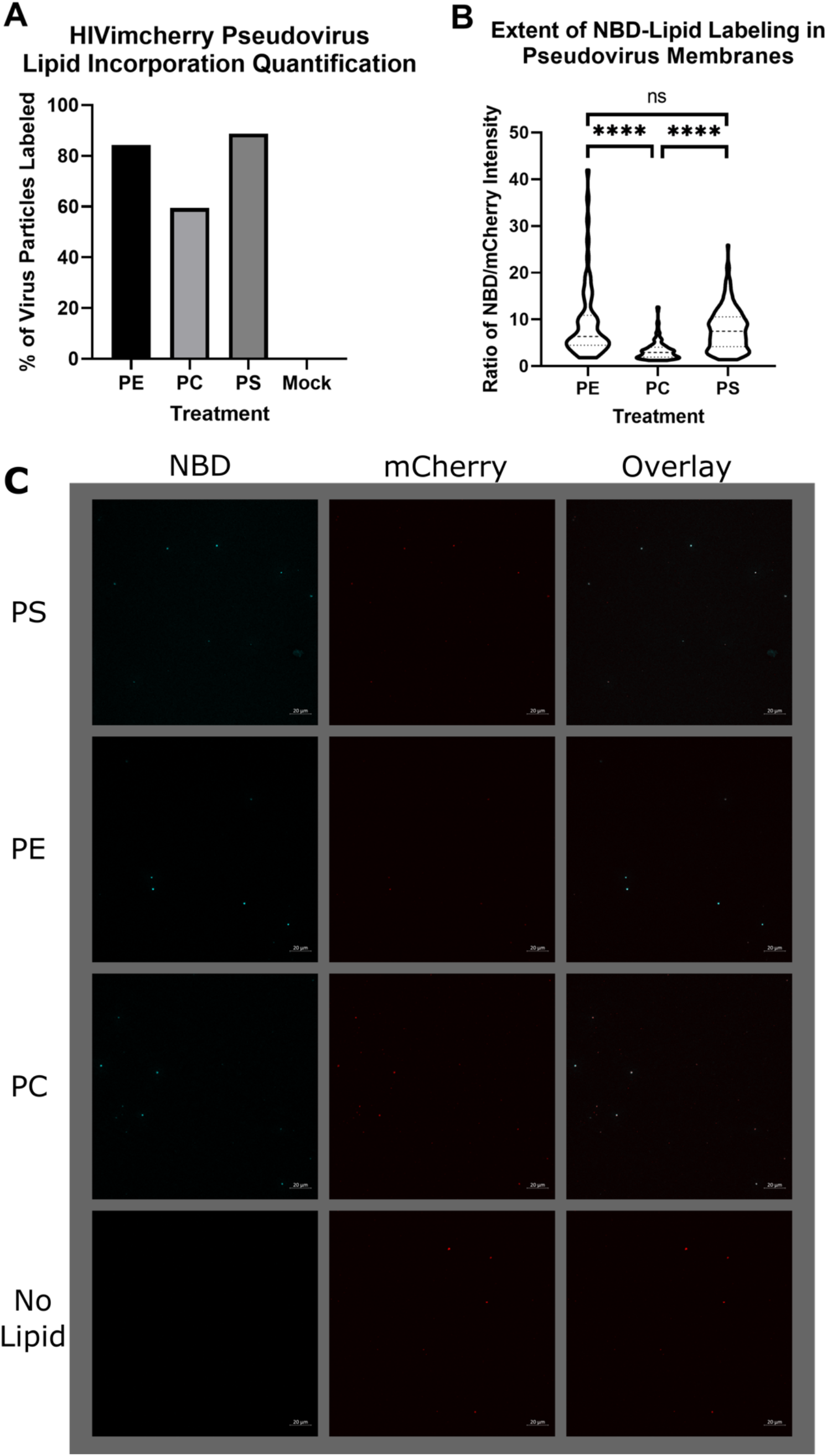
Incorporation of exogenous lipids into HIV pseudoviruses. HIV pseudoviruses were treated with an NBD acyl chain-labeled version of the phospholipids listed. (**A**) The percentage of particles (defined as punctate mCherry fluorescent region of interest) exhibiting NBD fluorescence as visualized by confocal microscopy. (**B**) The ratio of NBD to mCherry fluorescence intensities of the particles that had incorporated NBD-lipid is plotted as a proxy for the relative amount of exogenous lipid incorporated into the viral membrane. The dashed line is the median and the dotted lines are quartiles. **** p<0.0001, ns not significant. (**C**) Representative fluorescence micrographs of the data quantified in (**A**) and (**B**). Each punctum is considered a single viral particle. NBD is pseudocolored in cyan and mCherry is pseudocolored red. Scale bars are 20 μm.

**Fig. S7.**
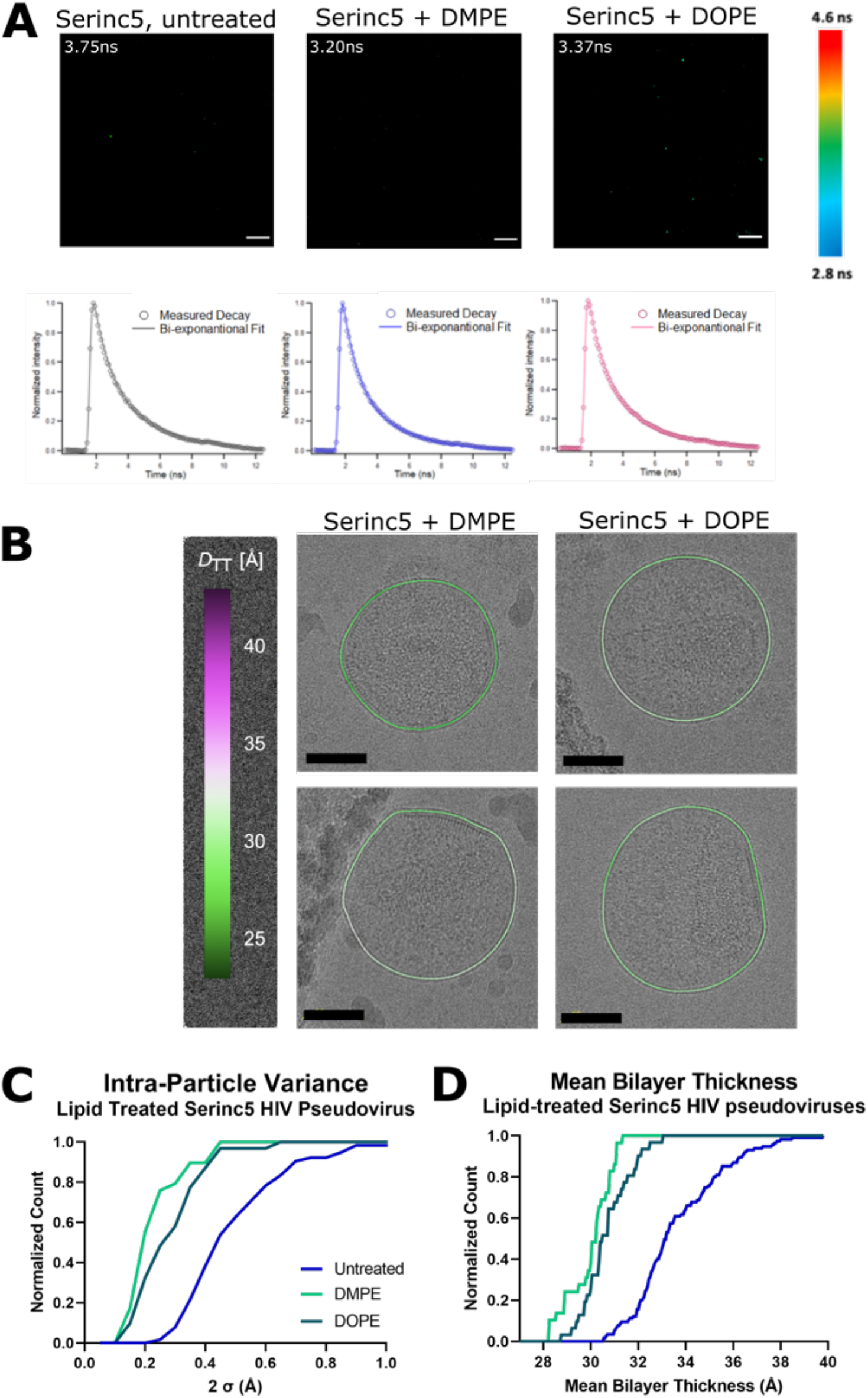
Effects of lipid enrichment of Serinc5-containing HIV pseudovirus particles on membrane order. (**A**) Example fluorescence micrographs of Serinc5-containing HIV pseudoviruses, enriched with the phospholipid listed above and then stained with FLIPPER-TR, adhered to a poly-L-lysine coverslip, and imaged. Lifetime is scaled in color as shown in the key to the right. The corresponding decay curves are shown below the micrographs for all pixels that exceed an intensity threshold to exclude background but include punctate fluorescence of virus particles. The calculated average lifetime from two component fit of the decay curve is written in white in the corner of the micrograph. Scale bars are 20 μm. (**B**) Example cryo electron micrographs of Serinc5-containing HIV pseudoviruses enriched with the exogenous phospholipids written above the column. Trough to trough distances (D_TT_) of viral membranes are shown as heat maps (scale on the right). All scale bars are 50 nm. (**C**) 95% confidence intervals of the mean bilayer thickness (D_TT_) of individual lipid-enriched HIV pseudovirus particles plotted as a cumulative distribution function. Unenriched Serinc5 curve replotted from Fig. 2B for comparison. (**D**) Mean membrane thickness (D_TT_) of individual lipid-enriched HIV pseudoviruses were plotted as a cumulative distribution function. Unenriched Serinc5 curve replotted from Fig. 2C for comparison.

**Fig. S8.**
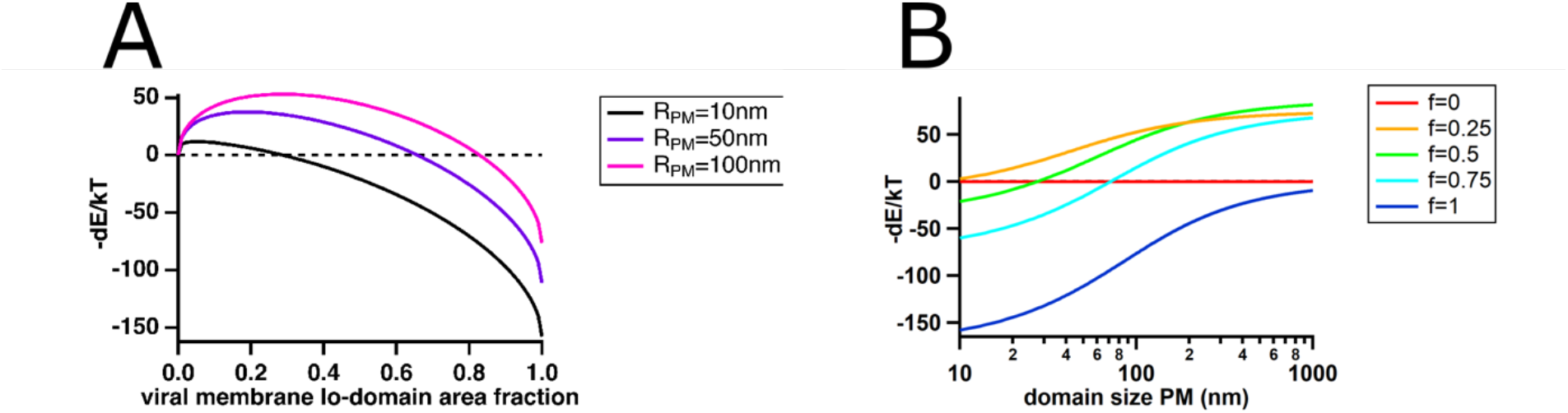
Contribution of line tension to energy change of fusion. Based on the model described in (Yang et al., 2016a) (see also Methods), the effect of increasing the fraction of the viral membrane in a single ordered domain on the contribution of line tension to the energy change of fusion is plotted in panel (**A**). The radius of the virus was held constant at 55 nm and the radius of domains in the plasma membrane were varied from 10 to 100 nm as shown in the key. Positive values of -dE/kT (above the dotted line) favor fusion. In panel (**B**), the effect of increasing domain fraction (f, in key) and size (x-axis) in the plasma membrane on the energy change of fusion is plotted. The radius and L_o_ domain area fraction of the viral membrane are held constant at 55 nm and 0.5, respectively. The calculations show that increasing the viral membrane fraction in the L_o_ phase has a negative effect on the fusion probability especially for domain sizes in the plasma membrane below 100 nm. If the whole viral membrane is in a L_o_ phase the free energy contribution will always be higher in the fusion product compared to the prefusion situation therefore making fusion less likely.

## Notes

### Competing Interest Statement

The authors have declared no competing interest.

